# Eradication of *ENO1*-deleted Glioblastoma through Collateral Lethality

**DOI:** 10.1101/331538

**Authors:** Yu-Hsi Lin, Nikunj Satani, Naima Hammoudi, Jeffrey J. Ackroyd, Sunada Khadka, Victoria C. Yan, Dimitra K. Georgiou, Yuting Sun, Rafal Zielinski, Theresa Tran, Susana Castro Pando, Xiaobo Wang, David Maxwell, Zhenghong Peng, Federica Pisaneschi, Pijus Mandal, Paul G. Leonard, Quanyu Xu, Qi Wu, Yongying Jiang, Barbara Czako, Zhijun Kang, John M. Asara, Waldemar Priebe, William Bornmann, Joseph R. Marszalek, Ronald A. DePinho, Florian L. Muller

## Abstract

Inhibiting glycolysis remains an aspirational approach for the treatment of cancer. We recently demonstrated that SF2312, a natural product phosphonate antibiotic, is a potent inhibitor of the glycolytic enzyme Enolase with potential utility for the collateral lethality-based treatment of Enolase-deficient glioblastoma (GBM). However, phosphonates are anionic at physiological pH, limiting cell and tissue permeability. Here, we show that addition of pivaloyloxymethyl (POM) groups to SF2312 (POMSF) dramatically increases potency, leading to inhibition of glycolysis and killing of *ENO1*-deleted glioma cells in the low nM range. But the utility of POMSF *in vivo* is dose-limited by severe hemolytic anemia. A derivative, POMHEX, shows equipotency to POMSF without inducing hemolytic anemia. POMHEX can eradicate intracranial orthotopic *ENO1*-deleted tumors, despite sub-optimal pharmacokinetic properties. Taken together, our data provide *in vivo* proof-of-principal for collateral lethality in precision oncology and showcase POMHEX as a useful molecule for the study of glycolysis in cancer metabolism.

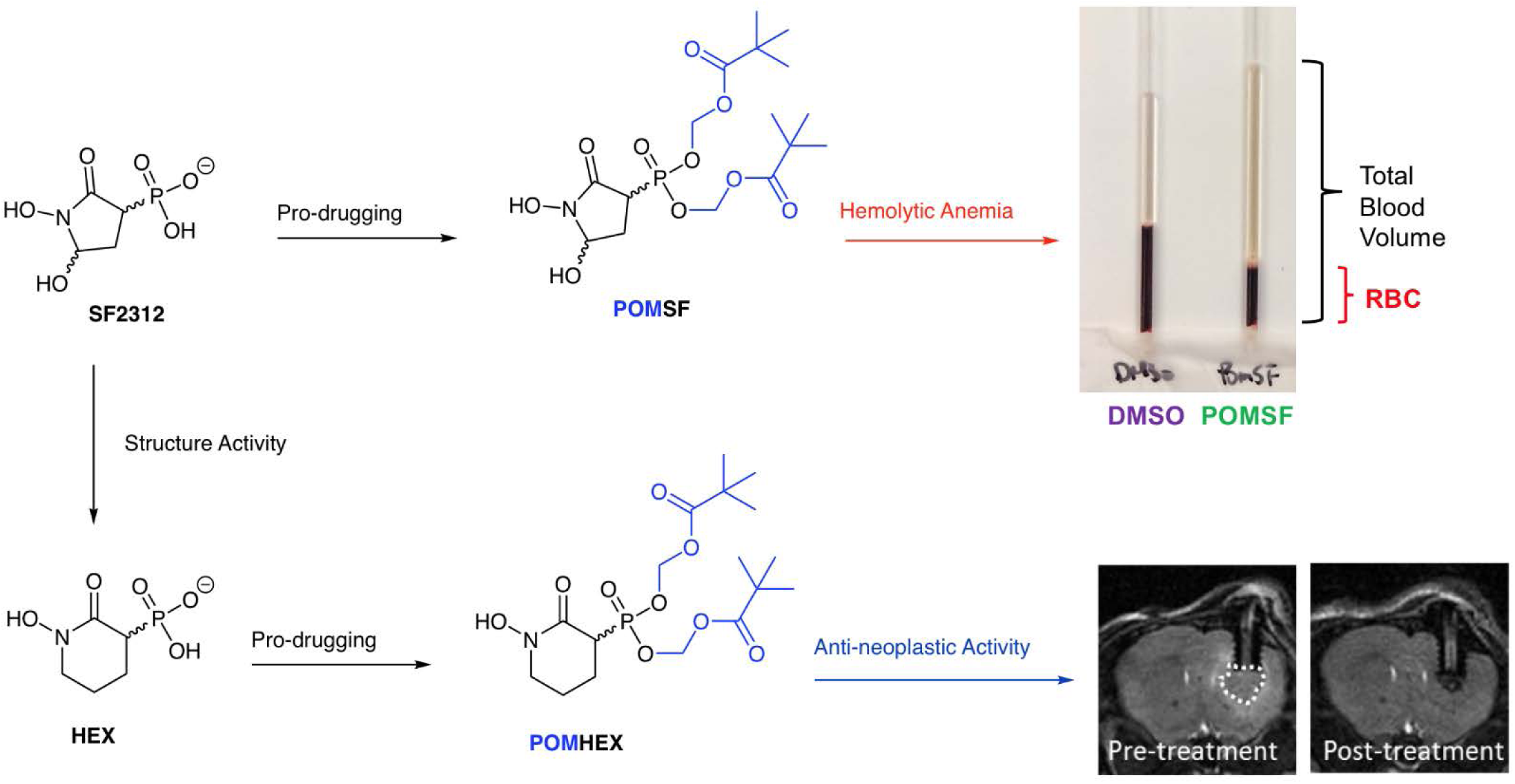

## INTRODUCTION

Glycolysis serves a critical role in cancer metabolism, as elevated glycolytic flux (lactate production) provides essential anabolic support for cellular growth and proliferation. Targeting glycolytic pathways has thus been an aspirational target for the treatment of cancer. However, few high-affinity inhibitors of glycolysis enzymes have been described, with most being tool compounds of limited utility beyond *in vitro* enzymology studies^1,2^. Yet the necessity of glycolysis for the viability of all cells—cancerous or not—begs the question as to whether a general inhibitor of glycolysis would show a sufficiently large therapeutic window for effective anti-neoplastic efficacy.

To explore this area, we conceived and validated the ‘collateral lethality’ strategy that takes advantage of passenger genomic deletions in tumor suppressor loci which create glycolytic vulnerabilities in cancer cells over normal tissue^3^. The first example of collateral lethality was accomplished by identifying tumors with homozygous deletions of the 1p36 tumor suppressor locus, which frequently incurs deletion of the glycolytic enzyme Enolase 1 (ENO1). Enolase activity is cell-essential and catalyzes the conversion of 2-phosphoglycerate (2-PG) to phosphoenolpyruvate (PEP) in the penultimate step of glycolysis. Tumors harboring homozygous deletions at this locus can still perform glycolysis through redundant action of Enolase 2 (ENO2).

Concurrent with this dependence on ENO2, we demonstrated that glioblastoma cells with homozygous deletion of ENO1 are exquisitely sensitive to shRNA mediated depletion of ENO2 ^3^. Though we initially thought that a selective ENO2 small molecule inhibitor would be required to reproduce the effect observed by shRNA against ENO2, we discovered that a non-selective, pan-Enolase inhibitor could exhibit up to 50-fold selective toxicity against *ENO1*-null glioma cells compared to *ENO1*-intact glioma cells or normal human astrocytes^3^. This is because ENO1 is the major isoform of the enzyme; ENO1-homozygously deleted glioma cells only retain ∼10% of normal Enolase activity. That a pan-Enolase inhibitor is highly toxic to these cancer cells is a testament to their acute sensitivity, as only a small amount of inhibitor is required to inhibit the residual enzyme below toxic threshold.

We recently demonstrated that SF2312, a natural product phosphono-hydroxamate, is a potent inhibitor of the glycolytic enzyme Enolase that exhibits selective toxicity towards glioma cells with passenger homozygous deletions in ENO1^3^. While SF2312 would appear to be viable candidate for molecular targeted therapy, the efficacy of phosphonate-containing drugs in general is severely limited by their ionic state at neutral pH. This compromises meaningful cell and tissue permeability^3^. To alleviate this issue, we protected the charged phosphonates on SF2312 with the pivaloyloxymethyl (POM) moiety to generate POMSF (**1**). This dramatically increases the cell-based potency for inhibition of glycolysis and selective killing of *ENO1*-deleted glioma cells. At the same time however, *in vivo* administration of POMSF resulted in hemolytic anemia as a likely consequence of on-target activity against ENO1 in red blood cells, evidently undermining its utility.

To overcome this liability, we performed further structure-activity-relation (SAR) studies to identify our lead compound, POMHEX (**2**), which exhibits near equipotency with POMSF in cell-based assays while critically avoiding the issue of inducing hemolytic anemia—even at doses 20-fold higher. POMHEX (and the free phosphonate HEX **3**, at higher doses) clearly display anti-neoplastic activity against *ENO1*-null xenografted tumors in mice. Overall, these data provide *in vivo* evidence for collateral lethality as a viable strategy to target cancer-specific vulnerabilities arising from passenger deletions.

## RESULTS

### Improved cellular potency of SF2312 by POM modification

The POM pro-drug moiety is a well-established chemical modification used to increase cell permeability of ionic phosphonate-containing drugs by masking their negative charges^4,5^ (Figure 1a). These POM moieties are intracellularly removed through sequential action of carboxylesterase and phosphodiesterase—releasing the active phosphonate drug^4,5^. POM-group addition to SF2312 was thus a natural direction for us to examine both the effect of plasma membrane permeability on cell-based inhibition of glycolysis and selective toxicity to *ENO1*-null versus *ENO1*-intact glioma cells.

**Figure 1:**
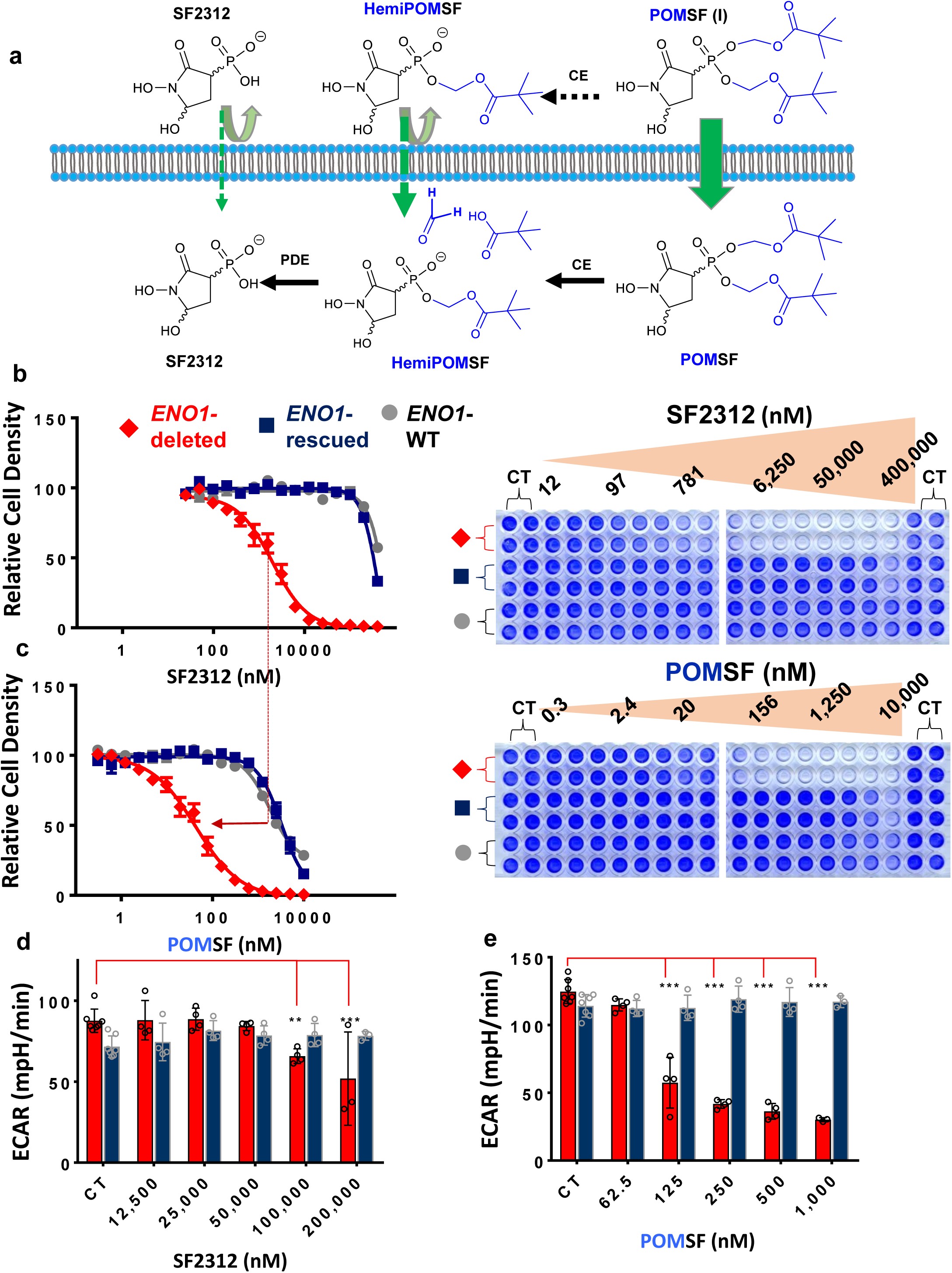
A POM pro-drug of SF2312 displays dramatically increased potency against glioma cells in vitro. **Panel a**: Negatively charged phosphonates like SF2312 have limited membrane permeability. POM groups (blue) mask these negative charges and enable membrane penetration. POM esters are removed intracellularly by the sequential action of carboxylesterases (CE) and phosphodiesterases (PDE)^5^. Carboxylesterase activity is also present in serum containing media, leading to slow hydrolysis (**Suppl Fig 1**). *ENO1*-deleted (D423, red), *ENO1*-isogenically rescued (D423 ENO1, blue), and *ENO1*-WT (LN319, grey) cells were treated for 7 days with SF2312 (b) and POMSF (c) at the doses indicated (nM). Cell density was quantified by crystal violet and expressed as a function of vehicle treated controls (CT). Addition of POM groups lowered the IC50s by ∼50-fold (red dashed line). Mean of N=20 biological replicates ± SEM are shown, with greater than three independent repetitions of the experiment (**Suppl. Table 1**). The effect of SF2312 (d) and POMSF (e) on glycolytic flux was quantified by extracellular acidification rates (ECAR) in *ENO1*-null (D423, red bars) and isogenic *ENO1*-rescued cells (D423 ENO1, blue bars). Individual data points and the mean ± S.D. of N = 7 (CT) and N = 3 – 4 (SF2312, POMSF-treated) biological replicates are shown. Statistical analysis was done utilizing 2-way ANOVA with Tukey’s Multiple Comparison Test; significant differences are indicated \*\**P*<0.01,\*\*\**P*<0.001). One independent repetition of the experiment was performed.

The POM-adduct, POMSF (**1**), was synthesized from the benzyl-protected phosphonic acid precursor (**4**). Reaction with silver nitrate generated a silver salt, to which iodomethyl pivalate in toluene was added—yielding the benzyl-protected POM-adduct. Reductive hydrogenation for removal of the benzyl-protecting group produced the desired product (**Supplementary Note 1**). POMSF was stable in neutral phosphate buffer solutions for >7 days but exhibited a relatively rapid hydrolysis (half-life of <12 h) to the monoester form in cell culture media containing fetal bovine serum (FBS) (**Supplementary Figure S1**). This hydrolysis is most likely due to the presence of carboxylesterase in FBS^5,6^.

We systematically compared the effects of the free phosphonate SF2312 to its pro-drug derivate, POMSF, by analyzing its cell-based inhibition of glycolysis and selective toxicity to *ENO1*-null versus *ENO1*-intact glioma cells. These cell lines were previously described in detail^3,7^. Briefly, the D423-MG (H423 in^8^) glioma cell line is *ENO1*-null. Ectopic re-expression of ENO1 generates an isogenic clone (D423 ENO1). LN319 serves as an *ENO1*-intact glioma cell line control.

POMSF exhibits robust toxicity in *ENO1*-null glioma cells, with an IC_50_ of ∼50 nM. This is approximately 100-fold greater than the I_50_ of ∼2,000 nM for the parental free phosphonate in SF2312 (Figure 1b, c). But even this difference is likely an underestimate, owing to the relative instability of POMSF in cell culture media (**Supplementary Figure S1**). Importantly, POMSF retained the selective killing of *ENO1*-null over *ENO1*-intact glioma cells, whereas *ENO1*-isogenic rescued or otherwise *ENO1*-intact glioma cells were essentially unaffected at concentrations of POMSF less than 1,000 nM.

Consistent with its selective toxicity, POMSF also preferentially inhibits glycolysis in *ENO1*-null over isogenic rescued glioma cells. Measurement by Seahorse extracellular acidification rates (ECAR) showed statistically significant reductions in flux observed at concentrations as low as 62 nM (Figure 1d, e). By contrast, SF2312 requires addition in the high µM concentration range to achieve selective inhibition of glycolytic flux in *ENO1*-null glioma cells. Critically, these effects observed for POMSF were not due to cell killing, as viability was not affected during these short-term exposures (<5 hrs; (**Supplementary Figure S2**).

Next, we tested POMSF for safety and dose-limiting toxicity in healthy adult mice. A single tail-vein intravenous injection with as little as 1.5 mg/kg of POMSF in healthy, non-tumor-bearing nude mice resulted in a >50% drop in red blood cell fraction (hematocrit) which was also evident by the red-coloring of the plasma (Figure 2a, b, c). This hematocrit drop is most consistent with hemolytic anemia, though the precise etiology of this anemia is uncertain. In this regard, it is worth noting that a rare familial hemolytic anemia genetic disorder is associated with *ENO1* point mutations^9,10^ and red blood cells are also exceptionally sensitive to glycolysis inhibition as they depend entirely on glycolysis for ATP generation due to an inherent lack of mitochondria^11^. While hemolytic anemia was fully reversible upon POMSF discontinuation (Figure 2a), this pathology severely limits the dose of POMSF that can be safely administered and be used for treatment of *ENO1*-deleted tumors. We thus sought to overcome this liability by synthesizing further derivatives.

**Figure 2:**
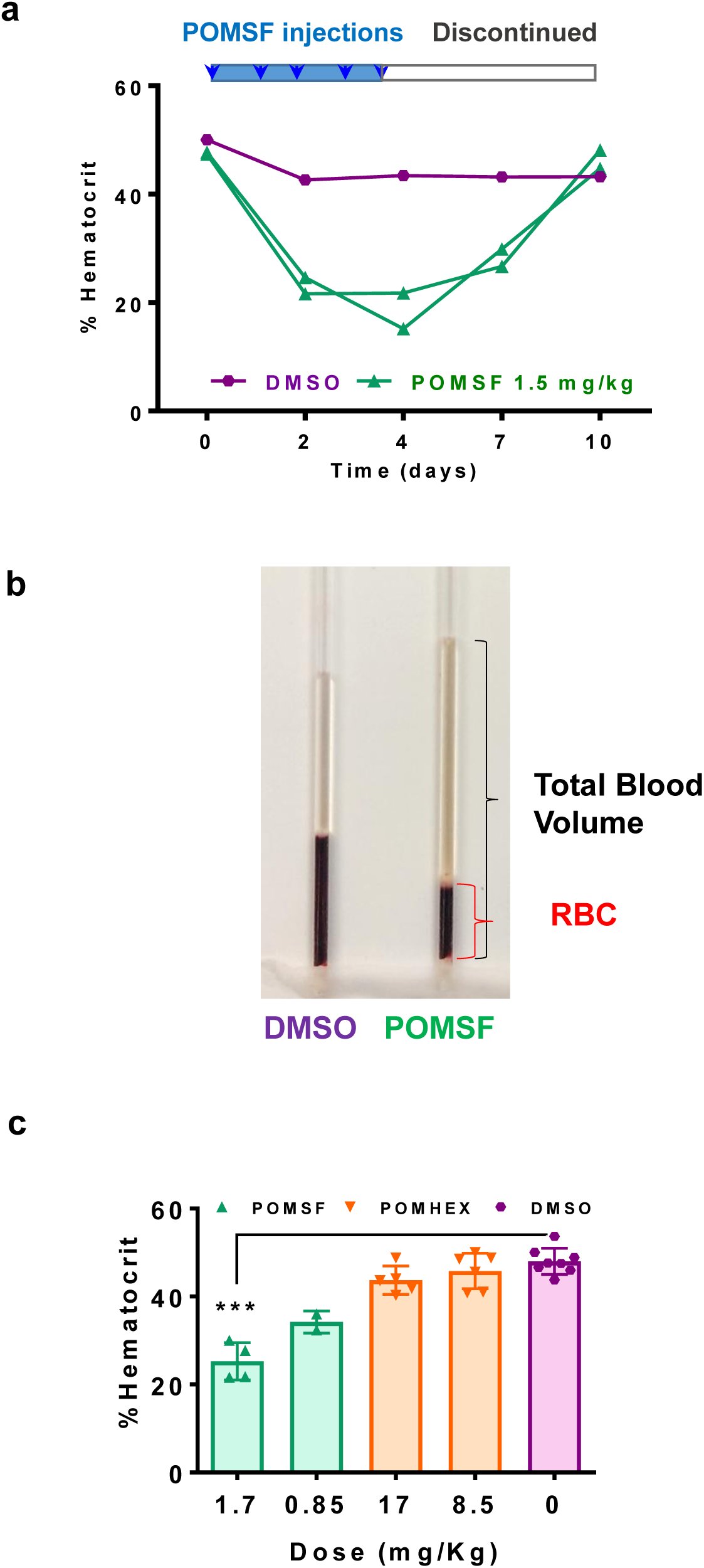
POMSF but not POMHEX causes hemolytic anemia in vivo. **Panel a**: Hematocrit in mice treated according to the indicated time schedule via intravenous injection with DMSO vehicle control (purple) or POMSF (green, 1.5 mg/kg) as a function of time. Each trace represents a single animal treated and sampled over consecutive days. **Panel b:** Representative capillary tube after centrifugation. The ratio of red blood cell fraction (RBC) to total blood volume is decreased in response to POMSF treatment. Additionally, yellowish plasma indicates hemolysis. **Panel c**: Hematocrit was determined after one week of intravenous daily administration of POMSF (green, N= 4, 2), POMHEX (beige, N = 5, 6), or DMSO vehicle treatment (pink, N = 8) at the doses indicated. Each data point represents one animal with mean ± S.D. POMHEX did not decrease hematocrit even when administered at doses 10-times greater than POMSF.

### POMHEX is near equipotent to POMSF in cell-based systems and does not cause hemolytic anemia

Because red blood cells possess only ENO1 and not ENO2, we reasoned that increasing the specificity of an Enolase inhibitor for ENO2 over ENO1 could mitigate the issue of hemolytic anemia. The IC_50_ of SF2312 for the inhibition of ENO1 and ENO2 are nearly identical despite the divergence in IC_75_^7^. To identify modifications that would improve target specificity for ENO2, we conducted SAR studies. Expanding the ring size from 5 to 6 atoms generated HEX (**3**, **Supplementary note 2**, Figure 3a), a substrate-competitive inhibitor with a clear preference for ENO2 over ENO1.

**Figure 3:**
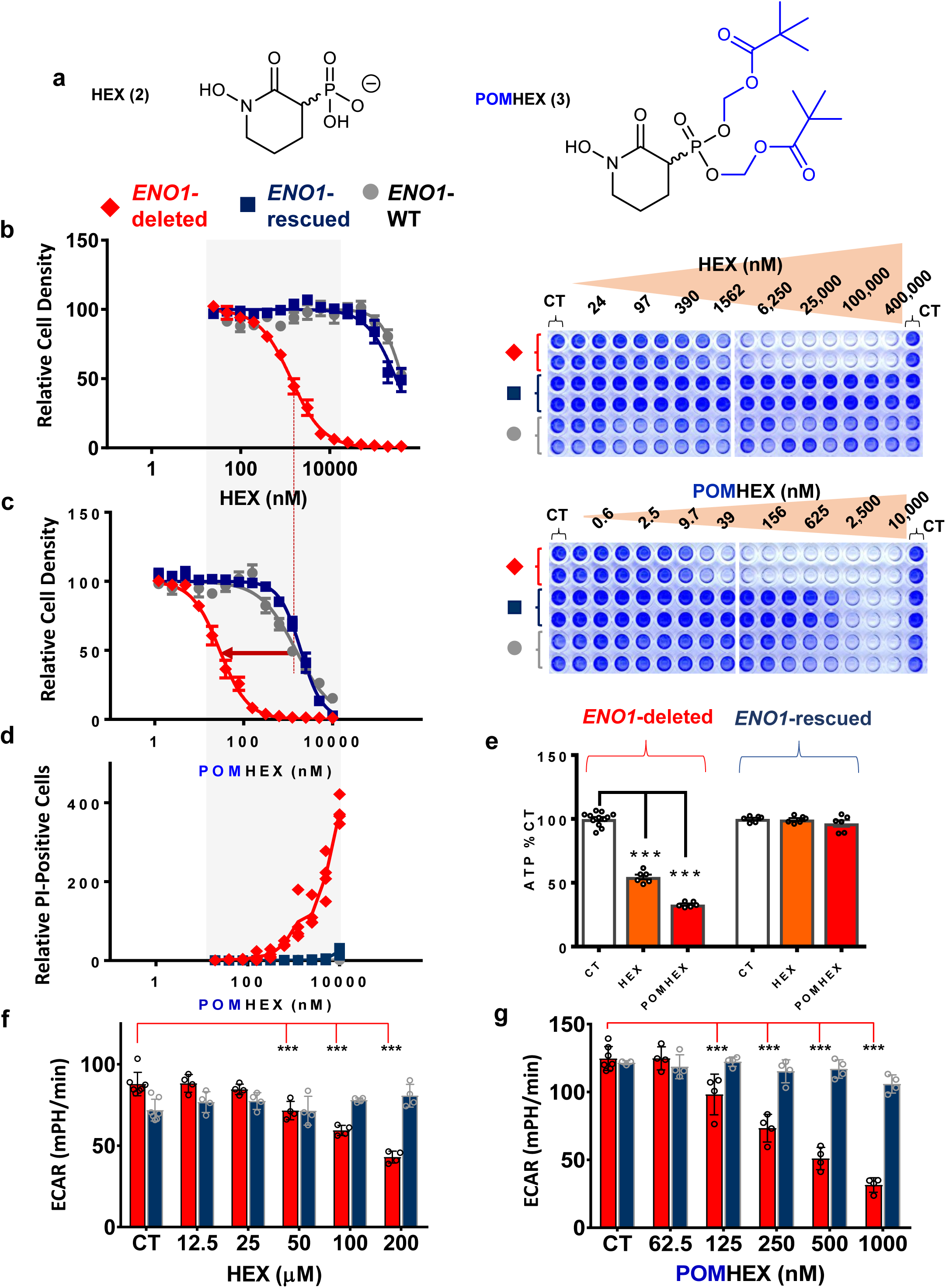
POMHEX selectively inhibits glycolysis and kills ENO1-null glioma cells. **Panel a**: Structure of the Enolase inhibitor HEX and its pro-drug derivative POMHEX with POM groups shown in blue. **Panel b,c**: *ENO1*-null (D423, red), *ENO1*-isogenically rescued (D423 ENO1, blue), and *ENO1*-WT (LN319, grey) cells were treated for 7 days with HEX (b) and POMHEX (c) at the doses indicated (nM). Terminal cell density was quantified with crystal violet (representative plates shown) and expressed as function of vehicle-treated controls (CT). Each data point represents the mean ± S.E.M. of N = 20 biological replicates. **Panel d:** POMHEX selectively induces cell death. Each data point represents a single biological replicate (N = 3-4 per concentration) of Propidium iodide positive cells relative to CT in *ENO1*-null (D423, red), *ENO1*-isogenically rescued (D423 ENO1, blue), and *ENO1*-WT (LN319, grey) cells. **Panel e:** D423 *ENO1*-null and D423 *ENO1*-rescued glioma cells were treated with HEX (orange bars, 200 μM) and POMHEX (red bars, 78 nM). Cells were treated for 8 hours and ATP was measured with the cell titer glow assay. Individual data points and the mean ± S.E.M. of N = 12 (CT) and N = 6 (HEX, POMHEX) biological replicates are shown. Statistical analysis was done using 1-way ANOVA with Tukey’s multiple comparisons test. Significant differences are indicated, \*\*\**P*<0.001. **Panel e,f**: The effect of HEX (f) and POMHEX (g) on glycolytic flux was quantified by extracellular acidification rates (ECAR) in *ENO1*-null (D423, red bars) and isogenic *ENO1*-rescued cells (D423 ENO1, blue bars). Individual data points and the mean ± S.D. of N = 7 (CT) and N = 4 (HEX, POMHEX treated) biological replicates are shown. Statistical analysis was done using 2-way ANOVA with Tukey’s Multiple Comparison Test. Significant differences are indicated, \*\*\**P*<0.001.

Co-crystallization of ENO2 and HEX (**Supplementary Figure S3, Supplementary Table S1, PDB: 5IDZ;** see legend in **Table S1** for more details) allowed us to identify key residue interactions in the active site. HEX binds to ENO2 by chelating the hydroxamate moiety to the Mg^2+^ cation and by forming a salt bridge between the anionic phosphonate and the cationic arginine 373 residue (**Supplementary Figure S3b, c**). Comparing the co-crystal structures of ENO2:HEX and ENO2:SF2312 revealed the structural basis for the difference in inhibitory behavior between the two. Non-competitive inhibition by SF2312 is explained by the strong interactions of the 5’ hydroxyl with the catalytic backbone residues of ENO2^7^, which locks the protein in a closed configuration. The absence of this 5’ hydroxyl in HEX keeps the active site in the open configuration without the second Mg^2+^ cation bound (**Supplementary Figure S3, S3**). In performing Michaelis-Menten titrations of the natural substrate (2-PG) and inhibitor (HEX), we found that HEX exhibits a *K*_i_ of 232 nM for ENO1 versus 64 nM for ENO2 (**Supplementary Figure S3**).

We then synthesized a POM derivative of HEX, POMHEX (**2, Supplementary note 2**) and compared the abilities of HEX and POMHEX to selectively kill *ENO1*-null glioma cells. During a one-week treatment course, HEX displayed an IC_50_ of approximately 1.3 µM against D423 *ENO1*-null glioma cells, a value comparable to that of SF2312 (compare Figure 3b versus Figure 1b). By contrast, the IC_50_ in non-target D423 *ENO1*-isogenic rescued and LN319 *ENO1*-WT glioma cells are >300 µM for both HEX and SF2312, highlighting a 200-fold target preference in glioma cells.

Concurrent with the trend observed for POM addition to SF2312, POM addition to HEX improved cell culture-based potency from the low µM to the low nM range (Figure 3b, c) while also displaying slightly greater stability in media than POMSF (**Supplementary Figure S1**). Treatment with POMHEX resulted in an IC_50_ of approximately 30 nM in D423 *ENO1*-deleted glioma cells compared to an IC_50_ of >1.5 µM for non-target *ENO1*-isogenic rescued glioma cells (D423 ENO1) or *ENO1*-intact (LN319).

We tested POMHEX against diverse cancer cell lines through submission to the NCI-60 (POMHEX NCI ID: NSC784584, **Supplementary Figure S4**) and Sanger Center cell line drug-sensitivity testing panels (POMHEX Drug ID #2148). On average, POMHEX is ∼50-fold more potent than HEX, though with substantial variation across cell lines (Range: 35-fold to 347-fold; **Supplementary Table S2**). We postulate this variation in sensitivity to POMHEX is contingent upon both *ENO1*-deletion status and the expression of carboxylesterases and phosphodiesterases (Figure 1a), which varies across cell lines. Our testing also confirmed that *ENO1*-null cells are the most sensitive POMHEX/HEX treatment. *ENO1*-heterozygous cell lines exhibit intermediate sensitivity (**Supplementary Table S2**), which aligns with our previous reports for phosphonoacetohydroxamate and SF2312^3,7^. Some groups of cell lines (e.g. melanoma in NCI-60 data set and B-cell leukemia/lymphoma in Sanger dataset) also exhibit unusual sensitivity. But no tested cell line even nears the sensitivity observed for *ENO1*-null lines.

To both cement the importance of hydroxamate chelation to the Mg^2+^ cation and honor the assiduous nature of the hydrogenation reaction in the final step of the synthesis, we tested BenzylPOMHEX (**Supplementary Note 1**)—the immediate precursor to POMHEX—for selective killing of *ENO1*-null glioma cells. In the absence of a free anionic hydroxamate in BenzylPOMHEX, no selective toxicity towards *ENO1*-null cells was observed (**Supplementary Figure S5**). This coincides with our previous conclusion that a free hydroxamate is required for Enolase inhibition^7,12^ and suggests that benzyl ethers cannot be removed in glioma cells.

Degradation of POMHEX to its hemi-ester form in cell culture media (**Supplementary Figure 1b, d**) prompted our investigation into determining whether this intermediate, HemiPOMHEX, could retain selective killing of *ENO1*-null glioma cells. We found that HemiPOMHEX was about 4-times more potent than HEX but much less potent than POMHEX in killing *ENO1*-null glioma cells (**Supplementary Figure S6**). This is consistent with previous studies on POM-protected phosphonates, which suggests that mono-POM ester phosphonates enjoy a mild permeability advantage over their non-esterified counterparts^4^.

Because intact POMHEX is almost entirely degraded by 24 hrs in cell culture media, we performed pulsed experiments: glioma cells were exposed to POMHEX for 1 hr, after which non-drug media was put in place. We found that, while 1 hr pulsed POMHEX yielded nearly identical levels of killing in *ENO1*-null glioma cells, these conditions resulted in considerably less killing of the *ENO1*-intact glioma cell lines (**Supplementary Figure S7**). Synonymously, a wider therapeutic window can be achieved with pulsed—rather than continuous—exposure. This is perhaps because *ENO1*-intact glioma cells can recover more quickly from Enolase inhibition. Broadly, this finding has implications for pharmacology and for the interpretation of treatment experiments with POMHEX *in vivo*. With this favorable result from the pulsed experiment in mind, we propound the following model of bioactivation. An initially uncharged POMHEX rapidly diffuses into cells—first hydrolyzed by high intracellular carboxylesterase activity and thus trapped in the negatively charged mono-ester form. Then slowly, through the action of phosphodiesterases^4^, the mono-ester form is converted to the fully active Enolase inhibitor, HEX.

We quantified the selective toxicity of POMHEX using cell death markers (propidium iodide), which confirm both the strong killing by POMHEX against *ENO1*-null glioma cells versus non-target control cells (Figure 3d) and a selective depletion of ATP (Figure 3e). POMHEX also preferentially inhibits glycolysis (ECAR, Figure 3f, g) in *ENO1*-null over isogenic rescued glioma cells as measured by Seahorse extracellular acidification rates (ECAR, Figure 3f,g). For D423 *ENO1*-null glioma cells, POMHEX inhibits glycolysis in the nM range whereas *ENO1*-intact cells require concentrations >1 µM. These effects were not due to fewer cells, as the number of viable cells remained unchanged during these short-term exposures to POMHEX (<5 hrs; **Supplementary Figure S2**).

Together, our data indicate that POMHEX/HEX are broadly equivalent to POMSF/SF2312 in cell-based systems for inhibiting Enolase, blocking glycolysis and selectively killing *ENO1*-deleted glioma cells. *In vivo* mouse experiments show that, unlike POMSF, POMHEX intravenous injections of up to 17 mg/kg/day for 1 week does not lead to anemia in mice (Figure 2e). We conclude that POMHEX possesses clear advantages over POMSF for anti-neoplastic *in vivo* experiments.

### Inhibition of Enolase exerts far-upstream disruptions in the hexose amine biosynthesis pathway and the non-oxidative pentose phosphate shunt

POMHEX is a highly useful tool compound for studying the biochemical consequences of glycolysis inhibition. *ENO1*-null, *ENO1*-rescued and *ENO1*-intact glioma cells were exposed to increasing doses of POMHEX (15 nM to 720 nM) over a short time course (72 hrs). These concentrations of POMHEX have minimal consequences on *ENO1*-rescued/intact glioma cells but profoundly affect *ENO1*-null glioma cells. Analysis of the effects of POMHEX on protein cell signaling markers (Figure 4a) for cell proliferation (phosphor-Histone H3, PLK1), cell death (Cleaved caspase 3), and stress (p-NDRG1, p-JunB) revealed three salient points. First, even at the highest POMHEX concentrations used (720 nM), none of these markers were altered in *ENO1*-rescued or *ENO1*-intact glioma cells. Second, in *ENO1*-null cells, we find that proliferation is inhibited with the lowest levels of POMHEX (>15 nM, reduction p-Histone H3), whilst induction of apoptosis (cleaved caspase 3) occurs at higher concentrations (>180 nM). Third, a stress response (elevated p-NDRG1, p-Jun) is evident at ∼180 nM.

**Figure 4:**
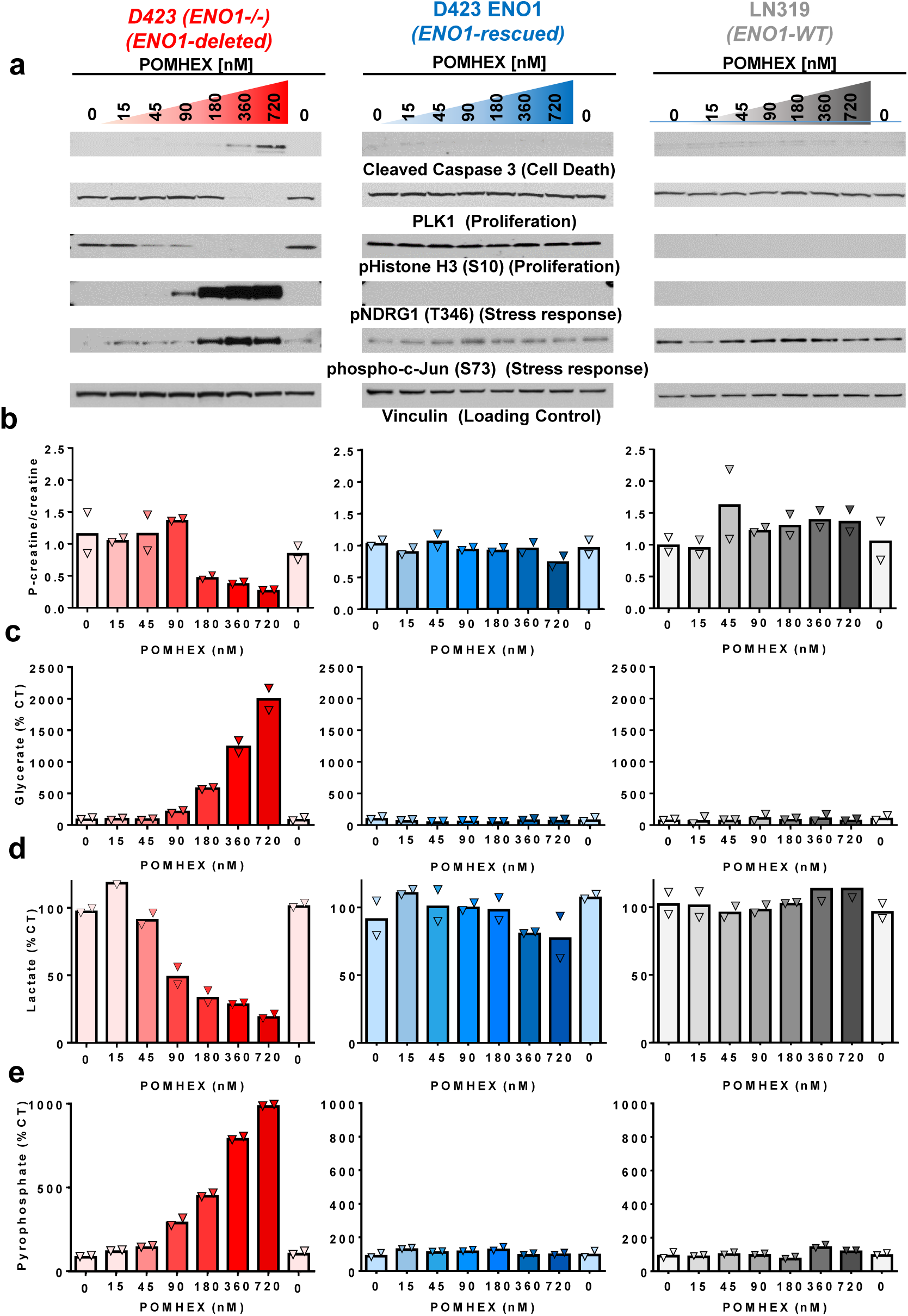
The Enolase inhibitor POMHEX selectively induces energy stress, inhibits proliferation, and triggers apoptosis in ENO1-null glioma cells. Glioma cells that are *ENO1*-null (D423, red), *ENO1*-isogenically rescued (D423 ENO1, blue), or *ENO1*-WT (LN319, grey) were treated with a dose response of the Enolase inhibitor POMHEX for 72 hrs before being harvested for protein lysates or polar metabolites. **Panel a:** *ENO1*-null cells experience a POMHEX dose-dependent increase in markers of the stress response (phosphorylation of T346 NDRG1, S73 c-Jun), decreased proliferation (phosphorylated Histone H3, PLK1), and increased cell death (cleaved Caspase 3) as indicated by immunoblot. These effects are specific for *ENO1*-null cells (left panel) and are not observed in *ENO1*-rescued or *ENO1*-WT cell glioma cells under the same treatment (right panels). Uncropped blots are shown in Supplementary Figure S10. **Panel b:** Phosphocreatine to creatine ratio (y-axis) was measured by mass spec. under the exact same treatment conditions. The ratio of phosphocreatine to creatine showed a POMHEX dose-dependent decrease in *ENO1*-null but not *ENO1*-rescued glioma cells [N = 4(CT), mean of N=2 (treated biological replicates)]. **Panel c:** Glycerate levels were measured as a proxy for blockage of the Enolase reaction by mass spec. in treated glioma cells and expressed as % of CT [(N = 4(CT), mean of N=2 (treated biological replicates)]. A dose-dependent increase in glycerate levels was observed in *ENO1*-null but not *ENO1*-rescued glioma cells. **Panel d:** Lactate levels were measured by ^1^H NMR with the integral of 1.34 ppm doublet normalized to 3-(trimethylsilyl)propionic-2,2,3,3-d4 acid standard and expressed as % of CT [(N = 2(CT), mean of N=1 (treated biological replicates)]. Note the dose dependent decrease in lactate levels in *ENO1*-null but not *ENO1*-rescued glioma cells. **Panel e:** Pyrophosphate levels were measured as a proxy for energy stress by mass spec in treated glioma cells and expressed as % of CT [(N = 4(CT), mean of N=2 (treated biological replicates)]. A dose-dependent increase in pyrophosphate levels was observed in *ENO1*-null but not *ENO1*-rescued glioma cells treated with POMHEX.

We correlated these cell-signaling markers to metabolite read outs of glycolytic flux (lactate production, Figure 4d) and Enolase inhibition (glycerate levels, Figure 4b), energy reserves (P-creatine to creatine ratio, Figure 4b) and energy stress (pyrophosphate, Figure 4e). Accordingly, stress response cell signaling biomarkers closely parallel the metabolites that indicate energy stress. Both p-NDRG1 and p-cJun become significantly elevated only at concentrations that simultaneously reduce the p-creatine/creatine ratio and increase the levels of pyrophosphate. In tandem, cell proliferation (P-histone H3) is inhibited at concentrations of POMHEX (45 nM) that are both too low to induce actual energy stress and only marginally reduce glycolytic flux (lactate secretion, Figure 4d). These data suggest that glioma cells have coping mechanisms that link reduced energy production (glycolytic flux) and with reduced energy expenditure (proliferation).

To understand the metabolic consequences and possible adaptations to inhibited glycolysis, we performed polar metabolite profiling on the same set of samples (dose response POMHEX treated *ENO1*-null/rescued glioma cells). The most significant metabolite alterations are shown in metabolic map summary format in Figure 5a, with the size of the up/down arrows proportional to the magnitude of alteration; the quantified metabolites plots are shown in Figure 5b. Chief amongst our findings is that inhibition of Enolase with POMHEX in *ENO1*-null glioma cells leads to a dramatic decrease in metabolites of the TCA cycle. For instance, we observed a >90% decrease in fumarate despite the pyruvate-supplemented cell culture media (1.25 mM, which is fact in far excess of the ∼70 µM on average, in human blood; human metabolome database, HMDB0000243). This suggests that, at least in this specific glioma cell line, glucose metabolized in glycolysis is a major source of anaplerotic substrates for the TCA cycle, with minor contributions from exogenous pyruvate or glutaminolysis uptake.

**Figure 5:**
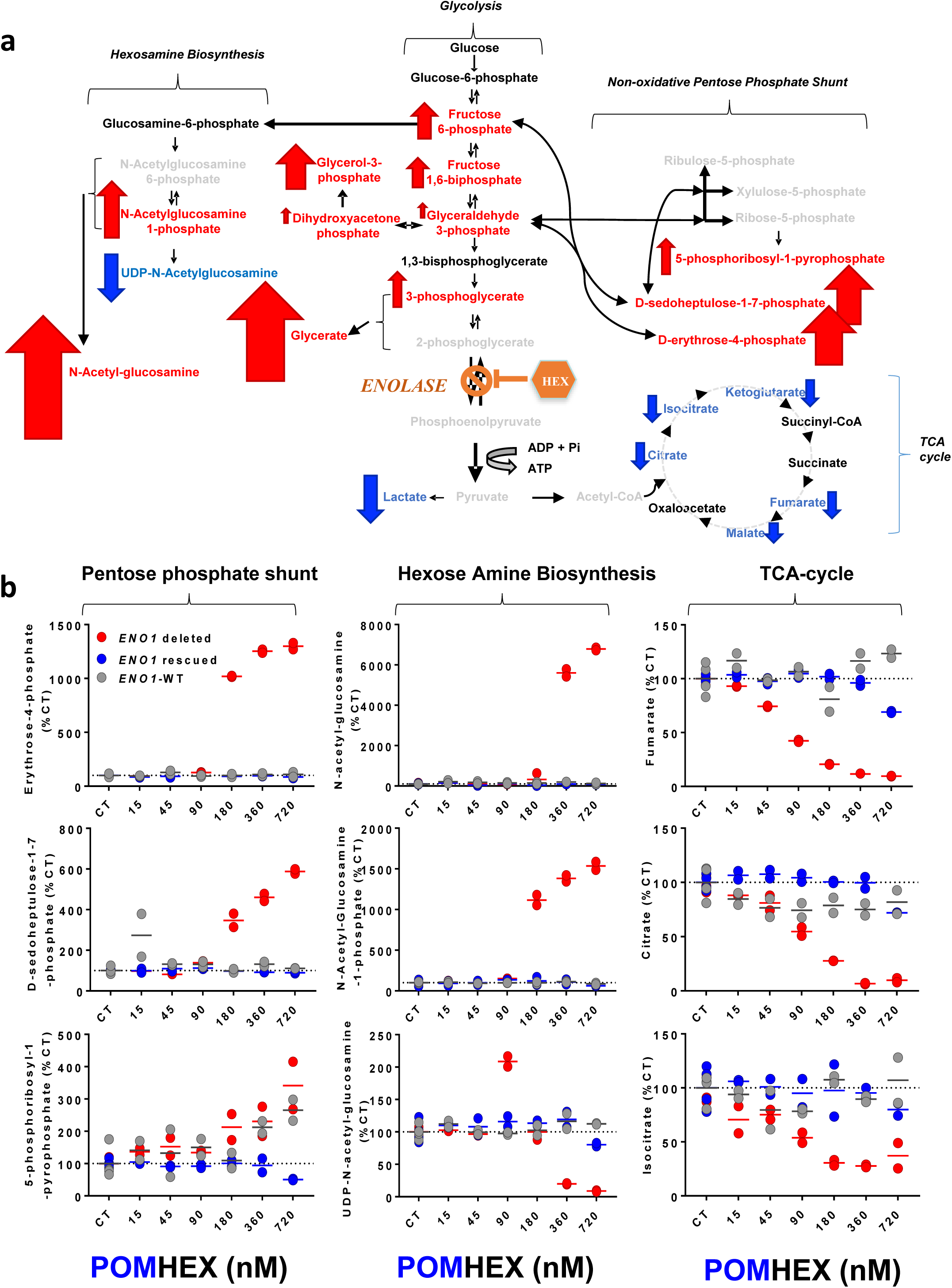
Inhibition of Enolase leads to spillover of metabolites into peripheral branches of the glycolysis pathway and depletion of TCA cycle intermediates. **Panel a**: graphical summary of metabolomic profiling of the effects of POMHEX treatment in *ENO1*-null versus *ENO1*-rescued/WT glioma cells. Red Arrows indicate metabolites that are selectively increased by POMHEX in *ENO1*-null versus *ENO1*-intact glioma cells, with the size of the arrow being proportional to the magnitude of the increase. Blue down-facing arrows indicate metabolites that are selectively decreased by POMHEX treatment in *ENO1*-null glioma cells. The main effects of inhibition of Enolase in *ENO1*-null glioma cells consists of an accumulation of metabolites at the periphery of glycolysis, resulting in disruptions in the pentose phosphate shunt and the hexosamine biosynthesis pathway. **Panel b:** Quantitative data of selected metabolites from panel a. *ENO1*-null (Red), rescued (blue), and intact (grey) glioma cells were treated for 72 hrs with POMHEX at the concentrations indicated (x-axis). Selected alterations from specific pathways are shown (N = 4 for CT, N = 2 for each concentration of POMHEX, independent biological determinations, with individual data points shown). Each metabolite is expressed relative to the mean of no-drug control (%CT). The experiment was performed once, as shown.

We thus repeated the POMHEX sensitivity experiments while titrating the pyruvate concentration in cell culture media. Omitting pyruvate did sensitize glioma cells to POMHEX— decreasing the IC_50_ of D423 *ENO1*-null cells by about 2.5-fold compared to regular DMEM (used throughout the rest of the manuscript), which contains 1.25 mM pyruvate. Supplementing the media with even more pyruvate (5 mM) made only a marginal difference (**Supplementary Figure S8**). These relatively modest shifts in IC_50_ lead us to conclude that the loss of anaplerotic substrates from inhibiting glucose-derived pyruvate is a minor contributor to the toxicity of Enolase inhibition. Instead, we conclude that most of the toxicity is due to bioenergetic failure derived from lost glycolytic ATP production (Figure 3e, Figure 4b, e).

While we initially expected that the metabolites immediately upstream (2-PG, 3-PG, glyceraldehyde 3-phosphate) of the Enolase reaction would experience the greatest alterations, we found that the most significant metabolite accumulations occurred at the peripheral arms of glycolysis—specifically, the hexose amine biosynthesis pathway (HBP) and the non-oxidative pentose phosphate shunt (Figure 5a, b). Prominent amongst the HBP metabolite buildups were a >15-fold accumulation of N-acetyl-glucosamine-phosphate and >60-fold accumulation of its hydrolysis product, N-Acetyl-glucosamine. The pattern of metabolite alterations in the HBP defies simple interpretation—as exemplified by a >90% decrease in the final HBP metabolite, UDP-N-acetyl-glucosamine, in response to POMHEX. We propose that excess metabolic flux from glucose “overflows” into these peripheral carbohydrate metabolic pathways in response to the blocked central flux at the Enolase step in glycolysis. Subsequent disruptions in the HBP could result in aberrant glycosylation, as HBP metabolites are the main substrates for post-translational protein glycosylation. This is reinforced by previous studies which tie alterations to HBP metabolites with perturbed patterns of protein glycosylation^13^.

In addition to the HBP, the non-oxidative pentose phosphate shunt^14^ is also disrupted in response to Enolase inhibition. We observed up to 12-fold elevations in D-sedoheptulose-1,7-phosphate and erythrose-4-phosphate: two metabolites that are interconverted with the main sequence glycolytic metabolites, fructose-6-phosphate (elevated ∼6-fold by POMHEX, Figure 5b) and glyceraldehyde-3-phosphate^14^ through action of transaldolases and transketolase enzymes. The functional consequences of D-sedoheptulose-1,7-phosphate and erythrose-4-phosphate accumulations are presently unclear. However, it is tempting to attribute the disruption of the normal flux through the non-oxidative pentose phosphate shunt to this metabolite imbalance and the shifted equilibrium of the transaldolase and transketolase reactions which follow. Should this hold true, redox balance and nucleotide pools would be affected—compromising cell viability and likely contributing to the toxicity of Enolase inhibition.

### POMHEX and HEX display anti-neoplastic activity against *ENO1*-deleted orthotopic tumors

Next, we performed safety and toxicity testing with HEX and POMHEX in mice, using the same strain of Nude mice that we would use for anti-neoplastic studies. HEX was exceptionally well tolerated once it was neutralized with NaOH to generate the monosodium salt: intravenous injections at 150 mg/kg/day are tolerated for months without overt signs of toxicity such as hemolytic anemia or loss of body weight. Without neutralization however, HEX is only tolerated with intravenous injections of up to 75 mg/kg most likely due to tissue and blood acidification.

We note the great technical difficulty in repeatedly punching the tail vein of mice for intravenous injections. Mice can tolerate intravenous injections of up to 300 mg/kg/day. HEX administered at 300 mg/kg to a mouse of 30 g with a plasma volume of ∼2 mL, would result in a C_max_ of nearly 20 mM HEX (compare to ∼5 mM plasma phosphate). Irrespective of its effects on Enolase activity, such concentrations of HEX will inevitably alter blood chemistry and are bound to be acutely toxic. Thus, we propose that even higher doses could either be administered using slow infusion pumps or punching the tail vein multiple times per day to limit the C_max_ of HEX below 5 mM in plasma. We also tried sub-cutaneous injections and found that up to 600 mg/kg, twice per day are tolerated without loss of body weight or anemia. Doses of 1200 mg/kg twice per day are also tolerated, but not without body weight loss and anemia (though both are fully reversible).

The toxicological profile of POMHEX was quite different. Intravenous injections were tolerated chronically at 10 mg/kg/day, with the maximum single tolerated dose at 30 mg/kg. Even at these relatively high doses, hemolytic anemia was not observed. Instead, we observed loss of body weight and sub-cutaneous fat—though both are reversible upon drug discontinuation.

Having established the toxicological profiles of both HEX and POMHEX, we then tested their anti-neoplastic effects in intracranial orthotopic *ENO1*-deleted tumors generated from the same D423 glioblastoma cell line we have utilized for *in vitro* experiments. D423 *ENO1*-deleted glioma cells were implanted in the left ventricle area of Nude (*Foxn1*^nu/nu^) mice^15^; tumors typically become T2-MRI detectable around 20-30 days post-implantation. At this point, tumor bearing mice were sorted by tumor size and consecutively assigned to drug treatment or no-drug CT arms such that tumor volume was the same in both groups. Animals were treated for a week with daily intravenous and intraperitoneal injection of either HEX (150 mg/kg in PBS) or POMHEX (10 mg/kg in PBS). After 1 week of treatment, T2-MRI was repeated and we determined the effect of treatment on tumor volume changes (Figure 6a). Animals were then sacrificed for biochemical profiling of pharmacodynamic target engagement markers.

**Figure 6:**
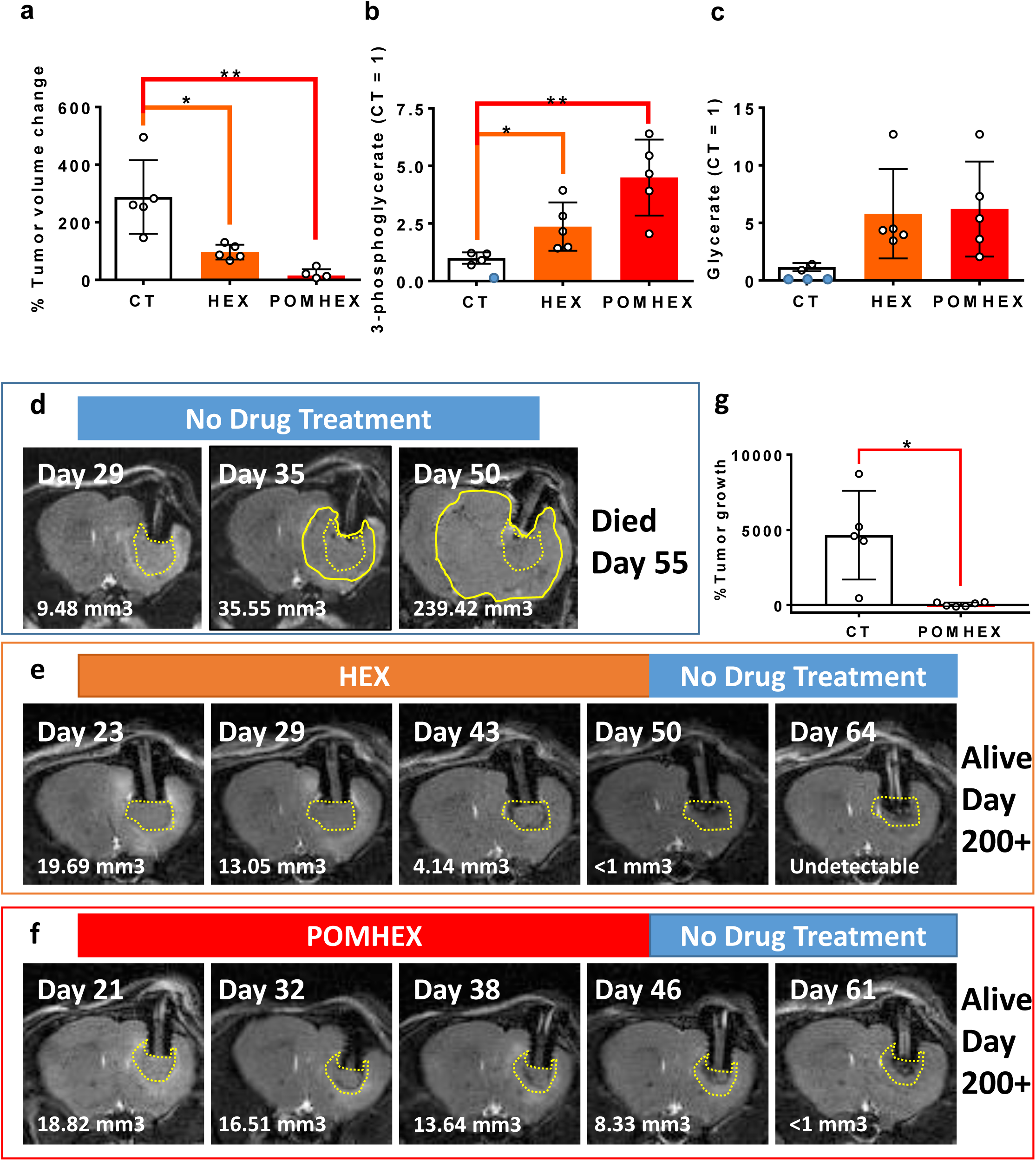
Anti-neoplastic and pharmacodynamic effects of Enolase inhibitors in orthotopic tumors. Intracranial tumors were generated by implantation of D423 (*ENO1*-null) glioma cells in nude immunocompromised mice and tumor formation was followed by T2-MRI. **Panel a** – tumor volume changes for untreated, HEX (150 mpk IV + 150 mpk IP), POMHEX (10 mpk IV + 10 mpk IP) tumor bearing mice (Mean of N = 5 +/− S.D. with individual data points shown) over the course of 1 week; after which animals were sacrificed and Enolase inhibition pharmacodynamic read-out metabolites were measured in tumors, including 3-phosphoglycerate (**b**) and glycerate (**c**); mean of 5 biological replicates +/− S.D, blue data points indicate readings below detection limit. Panel d,e,f: long term treatments; **Panel d**: Tumors become MRI detectable (dashed yellow outline) around 20-30 days after implantation; the number of days post-implantation is shown in upper left; tumor volumes were calculated from stacked images (in mm3 in each image); tumor volume increases inexorably in the absence of treatment, leading to infiltration of the entire brain and death of the animal. **Panel e**: Treatment with the non-prodrug HEX (75 mg/kg IV and 75 mg/kg IP) was initiated at day 23 post-implantation until it had regressed to the point of negligibility (day 50), at which point drug treatment was discontinued. The animals were effectively cured as the tumors did not recur even after extended time after treatment discontinuation. **Panel f**: Tumor bearing animal was put on treatment with POMHEX (10 mg/kg IV + 10 mg/kg IP) and tumor volume changes followed as a function of time. **Panel g**: summary of tumor volume changes following 18 days treatment with POMHEX (N = 6 POMHEX treated tumor bearing animals, N = 5 non-drug treated animals). (* P<0.05, **, P<0.01 by unpaired, 2-tailed, t-test with unequal variance using the Bonferroni correction). The experiments in d, e, f were repeated independently 2 times, while that in a, b, c were performed once, as shown.

Treatment by both POMHEX and HEX resulted in statistically significantly attenuated tumor growth compared to non-drug treated CT tumor bearing animals over the course of 1 week (Figure 6a, b). Pharmacodynamic target engagement markers provide confirmation of Enolase inhibition in intracranial tumors. As consistent with *in vitro* results, both HEX and POMHEX-treated intracranial tumors showed statistically significant elevations in glycerate and 3-PG compared to non-treated tumors (Figure 5). We also performed long-term (>2 weeks) treatment experiments, overcoming technical challenges such as multiple intracranial MRI imaging with long anesthesia and the fragility of the mouse tail vein during multiple punctures. Nevertheless, several longer-term treatment studies with either HEX or POMHEX showed eradication of intracranial *ENO1*-null tumors (Figure 6e, f). Strikingly, once tumors were eradicated, tumors did not recur upon treatment discontinuation. All treated tumor-bearing mice exhibited inhibition of tumor growth in response to Enolase inhibitors across all experiments conducted with the same protocol (Figure 6a). Complete eradication was observed in 6 out of 17 longer-term POMHEX treated mice and in 1 out 5 long term HEX treated mice.

We then examined *ex-vivo* and *in vivo* pharmacology of POMHEX and HEX. Consistent with the extensive literature on POM-phosphonate pro-drugs^5,6^, we found that POMHEX is very rapidly converted to the hemi-ester form, with a half-life of approximately 30 seconds (**Supplementary Figure S9**) in mouse plasma *ex-vivo*. However, the half-life in human blood *exvivo* was almost 9 minutes. This pronounced discrepancy in drug metabolism has previously been reported for other POM-containing phosphonate pro-drugs and is thought to be due to the significantly higher level of carboxylesterase activity in mouse plasma, compared to primates and humans^16-18^. Congruent with observations made for other POM-containing phosphonate drugs such as Adefovir and Tenofovir, POMHEX is expected to have a more favorable pharmacological profile in primates, including humans, compared to rodents. Pharmacokinetic analysis *in vivo*,following single IV or IP injections of POMHEX are in agreement with these ex-*vivo* results: intact POMHEX is essentially undetectable in mouse plasma, with only HemiPOMHEX and HEX being readily observed (**Supplementary Figure S9**).

We performed pharmacodynamic profiling in organs in mice treated for 1 week with IV/IP injections of POMHEX using the same protocol as for the anti-neoplastic experiments (Figure 6a). We observed that active HEX disproportionately accumulates in the heart (**Supplementary Figure S9d**), where the greatest degree of Enolase inhibition occurs, as evidenced by biochemical pharmacodynamic engagement markers (**Supplementary Figure S9,10**). Given the very short plasma half-life and the resulting steep concentration gradient away from the site of its injection in the venous system, this buildup in the heart is unsurprising. Both tail vein and IP-injections of POMHEX must traverse through the circulatory system before reaching the tumor (**Supplementary Figure S10**). Thus, only a small fraction of initially injected POMHEX ultimately reaches the tumor. Yet in spite of its poor pharmacology, POMHEX still cures intracranial tumors for two key reasons: 1.) *ENO1*-null glioma cells are at least 100-fold more sensitive to Enolase inhibitors, 2.) It is possible that the anti-neoplastic activity of POMHEX is at least partially mediated by its metabolites, HemiPOMHEX and HEX. HemiPOMHEX is more potent than HEX in cell-based systems (**Supplementary Figure 6**); HEX is active against *ENO1*-null intracranial tumors at higher concentrations (Figure 6).

In sharp contrast to the pharmacodynamic target engagement marker for POMHEX, mice treated with HEX at a dose 15-times higher gave much more favorable results. Lactate levels were not significantly decreased in any organ or in plasma while glycerate levels were only significantly increased in the brain (**Supplementary Figure S10**). Notably, *ENO1*-null brain tumors show a similar degree of elevation in glycerate and 3-phosphoglycerate as treatment with POMHEX (Figure 6c). Thus, treatment with HEX leads to a greater, on-target selectivity of Enolase inhibition in the brain and brain tumor, albeit requiring much higher concentrations than POMHEX.

It is worth clarifying how HEX might reach intracranial tumors, given its net negative charge and poor cell permeability. We propose two possible mechanisms. First, HEX may permeate intracranial tumors though a breached blood-brain barrier that is both characteristic of GBM and forms the basis for contrast-enhanced T1 MRI^19^. Alternatively, HEX may permeate the blood/cerebrospinal fluid barrier^20^, which is known to allow small highly polar drugs to pass. This includes the phosphonate antibiotic Fosfomycin^21^, a close structural analogue to HEX. Fosfomycin is present at a 1:10 plasma to cerebrospinal fluid barrier ratio^22^. Given the structural similarity between these two phosphonate compounds, we believe that HEX would act in a similar manner. That the normal, intact brain exhibits elevated pharmacodynamic markers of Enolase inhibition is compelling evidence that HEX can reach the brain and inhibit Enolase (**Supplementary Figure S10**).

## DISCUSSION

We previously demonstrated that SF2312, a natural product phosphono-hydroxmate, is an exceptionally potent Enolase inhibitor with selective toxicity against *ENO1*-null glioma cells^7^. However, a common drawback to all phosphonate-containing drugs, including SF2312, is poor permeability conferred by their net negative charge at physiological pH^26^. Of the approaches that exist to resolve this issue, we pursued pro-drugging for its synthetic simplicity and its well-established efficacy in the clinic, as evidenced by the highly successful anti-viral nucleotides HEPSERA and TRUVADA^26^. We thus generated POMSF.

Despite the enhanced permeability, the utility of POMSF ultimately suffers from inducing hemolytic anemia in mice, even at relatively low doses. Our subsequent efforts were thus devoted to perfecting the 6-membered ring, HEX, and its POM-adduct, POMHEX. Unlike substrate-noncompetitive SF2312, HEX is a substrate-competitive inhibitor with a distinct preference for ENO2. We believe that HEX does not induce hemolytic anemia (Figure 2) for two reasons: 1.) The exceptionally high levels of 2-PG in red blood cells hinder the inhibitory potency of substrate-competitive HEX but not of substrate-noncompetitive SF2312 and 2.) Red blood cells solely express ENO1.

We thus tested the anti-neoplastic effects of POMHEX in xenografted orthotopic intracranial *ENO1*-null glioma cells, the most faithful existing model for GBM pre-clinical drug testing, while also testing HEX as a control. POMHEX proved to be dramatically more effective than HEX on a molar basis for both inducing tumor regression and pharmacodynamic target engagement markers of Enolase inhibition in tumor and tissues (Figure 5, **Supplementary Figure 9**). Yet we were surprised by the efficacy of HEX in inhibiting Enolase and reducing tumor growth in the brain, despite its poor permeability. While it is near certain that negatively charged free phosphonates would not penetrate the blood brain barrier (BBB), small phosphonates with high structural similarity to HEX, such as Fosfomycin^30-34^and Foscarnet are extensively described in the literature as being able to readily penetrate the blood/cerebrospinal fluid barrier. Clinical effects in brain pathologies have been reported even in the absence of an obvious breach of the BBB^30-32,35-38^. HEX likely exerts an anti-neoplastic effect from its ability to penetrate the CSF/blood barrier.

While very little intact POMHEX reaches the brain and brain tumor due to its instability and steep concentration gradient in mouse plasma (**Supplementary Figure 10**), there is no obvious reason why intact POMHEX could not cross the BBB. Physiochemically, POMHEX is not too lipophilic (ADC lab predicted LogP: 0.76), has only one hydrogen bond donor, and has a molecular weight within the range where diffusion through the BBB is governed by lipophilicity^39^. It is also possible that, because POMHEX is quickly hydrolyzed to HemiPOMHEX and HEX in plasma, the combined penetration of these metabolites through the blood/CSF barrier contributes to the anti-neoplastic activity of POMHEX.

The poor pharmacology of POMHEX in mice (**Supplementary Figure S9,10**) could likely be rectified in human patients. Drug-metabolism of carboxylesterase-labile substrates differs significantly between rodents and primates. This is a well-recognized problem for using rodents as model systems for such drugs^16-18^. Carboxylesterase activity is about 80-fold higher in mouse compared to human plasma due to gene duplications restricted to the rodent lineage^18^, which protect against many toxins, including organophosphorus nerve agents^40^. However, this portends exceedingly short half-lives for phosphonate carboxylesterase-labile pro-drugs^5,41^. This evidently holds true for POMHEX, which would benefit from reduced carboxylesterase activity in human plasma to level its distribution across tissues and tumor.

This consideration, coupled with our pre-clinical data, strongly suggest that POMHEX would be effective against *ENO1*-null tumors in human patients, though the POM pro-drug is clearly not the most optimal pro-drug delivery system. Phosphonate/phosphate pro-drug development is an area of active research^26^. Pro-drug moieties with alternate bioactivation mechanisms have been synthesized. The use of a rodent model system warrants the exploration of pro-drugs with a non-carboxylesterase bioactivation to more readily assess the pharmacokinetics of future Enolase inhibitors. We believe an especially promising pro-drug strategy is based on bio-reduction, such as the nitrofurans pioneered by the Borch group^42^. This strategy exploits the low oxygen—and even hypoxic—tumor environment, which could result in a more favorable distribution of active Enolase in tumor versus normal tissues. Future work will be directed towards evaluating this and other moieties as an alternative pro-drug approach.

Finally, while the focus of this paper is on GBM, homozygous deletion of the 1p36 locus with *ENO1* also occurs in hepatocellular carcinoma, cholangiocarcinoma and large cell neuroendocrine lung tumors^25^—all cancers with poor prognoses and limited treatment options. Inhibiting Enolase targets the fundamental energy metabolism that all cells rely on to stay alive. Response to Enolase inhibition is thus solely contingent upon a tumor being *ENO1*-null. Indeed, at concentrations sufficient to inhibit ENO1 (10 µM) POMHEX is toxic even to *ENO1*-intact cancer cell lines, underscoring that glycolysis is universally necessary for cell viability. While the short-plasma half-life of POMHEX is problematic for systemic distribution, the drug would be ideally suited for intra-arterial interventional radiology treatment of tumors in the liver^44^. This would result in high levels of POMHEX close to the site of injection and swift diffusion into tissues/tumors. Rapid hydrolysis of POMHEX to HemiPOMHEX in the circulation ensures that systematic circulation would be exposed to poorly permeable HemiPOMHEX and HEX, mitigating toxicity to the rest of the organism. This same argument could be made for intra-carotid arterial delivery of POMHEX for GBM^45^.

In sum, for the first time, we demonstrate the efficacy of collateral lethality as a strategy for *in vivo*, pre-clinical treatment using a small molecule inhibitor. We clearly illustrate the robustness of this approach in our efforts to taking this concept one step closer to the clinic. Given the extensive catalogue of collaterally-deleted metabolic genes in cancer, collateral lethality stands to dramatically expand what may be considered actionable genetic alterations for precision oncology.

## AUTHOR CONTRIBUTIONS

F.L.M. and D.M. conceived HEX and POMHEX, which were synthesized by Z.P., D.S. and W.B.; Z.K., F.P., V.C.Y., F.L.M., P.M. and B.C. repeated chemical syntheses and wrote synthetic procedures with characterizations. N.S, and F.L.M. performed *in vitro* enzymatic activity inhibitor experiments. P.G.L. performed isolation of ENO2 recombinant protein and X-ray crystallography with HEX. R.Z. and W.P. performed Seahorse experiments. N.H., N.S., T.T., S.K., S.C.P., X.W. performed cell culture Enolase inhibitor sensitivity testing. J.J.A. performed western blots. Y-H.L. and F.L.M. performed ^13^C-NMR tracing and biochemical profiling experiments. J.M.A. performed Mass spec small molecule metabolite analysis. Q.X., Q. W., Y.J., Y.S. and J.R.M. performed pharmacological studies. X.W., S.K. and Y-H.L. performed mouse drug treatment and Y-H.L. performed small Animal MRI tumor imaging. J.R.M., Y.S., D.K.G. and F.L.M. generated figures, performed data analysis and statistics. F.L.M. and R.A.D. oversaw the project. Y-H.L., D.K.G., V.C.Y., N.S., R.A.D and F.L.M. wrote the manuscript.

## Methods

### Chemical synthesis and quality control

SF2312 was synthesized according to previously published procedures^7^. Initial syntheses of HEX and POMHEX were performed at M.D. Anderson’s Pharmaceutical Chemistry Facility. Subsequent syntheses were contracted to WuXi Apptec, Shanghai, China. Full synthetic descriptions are provided in Supplementary Note 1 (POMSF) and 2 (HEX/POMHEX). Product integrity was verified by ^1^H, ^31^P, ^13^C NMR and UPLC-MS in-house.

### Enolase enzymatic activity assay and inhibitor titrations

Enolase activity was measured *in vitro* using a plate reader as described both in previous publications^7,46^ and in the Nature Protocol Exchange (https://www.nature.com/protocolexchange/protocols/2430). For Michaelis-Menten titrations, the enolase substrate concentrations were titrated from 10 mM in 2-fold dilutions.

### Enolase inhibitor toxicity testing *in vitro* and cell lines

Cell culture experiments to test Enolase inhibitor sensitivity were conducted in 96-well plates. Plates were seeded at 1,500 cells per well. Cancer cells were attached for 24 hrs and were treated with fresh media containing Enolase inhibitor. Columns 1 and 2 as well as 11 and 12 were used as the media only CT. Columns 3-10 were used for drug treatment in 2-fold concentration gradients. After 6-7 days of growth, plates were washed with phosphate buffered saline (PBS) and fixed with 10% formalin. Fixed plates were stained with 0.1% crystal violet ^47,48^ and quantified by acetic acid extraction with spectrophotometric absorption at 595 nm in a plate reader. Cell densities were expressed relative to non-drug, media-only wells. Unless stated otherwise, all experiments were conducted in DMEM media with 4.5 g/L glucose, pyruvate and glutamine (Cellgro/Corning #10-013-CV) with 10% fetal bovine serum (Gibco/Life Technologies #16140-071) and 1% Pen Strep (Gibco/Life Technologies#15140-122) and 0.1% Amphotericin B (Gibco/Life Technologies#15290-018).

The main cell line used in this work is the H423/D423-MG (1p36 homozygous deleted, referred to as to D423 throughout the paper) and was kindly provided by D. Bigner^81^. The deletion in D423-MG spans the *CAMTA1, VAMP3, PER3, UTS2, TNFRSF9, PARK7, ERRFI1, SLC45A1, RERE, ENO1, CA6, SLC2A5, GPR157, MIR34A, H6PD, SPSB1*, and *SLC25A33* genes. An isogenic control cell line, D423 ENO1, where ENO1 is re-expressed ectopically, was described previously^3^. *ENO1*-heteroyzgously deleted glioma lines D502 (H502 in^8^) and U343-MG^49^ have also been described previously^3^. The ENO1-WT cell line LN319 is a sub-clone of LN-992,^50^though this detail is inconsequential for both the purposes described here and for the purposes of the CT in our experiments as any *ENO1*-intact line would be appropriate. The cell lines were authenticated and cultured as previously reported^7^.

### Western blots

Cell lysates were washed in ice cold phosphate-buffered saline (PBS). They were then collected in cold RIPA buffer with protease (Complete mini, Roche) and phosphatase inhibitors (PhosSTOP, Roche), and then sonicated. Protein concentration was determined by BCA assay (ThermoFisher 23227), separated by SDS-PAGE (4-12% gradiant), and transferred onto PVDF membranes using the Semi-Dry method (TransBlot turbo). Membranes were verified for efficient transfer with Ponceau S staining. Membranes were then blocked with 5% non-fat dry milk in TBS with 0.1% Tween 20 (TBST). Primary antibodies were incubated overnight at 4°C with gentle rocking. They were then washed 4x for 5 min with TBST. This was followed by secondary antibody incubation at room temperature for 1 hr with gentle rocking. Then, the membranes washed 4x for 5 min with TBST and incubated with ECL (ECL prime GE Healthcare (RPN2236)) or (ThermoScientific SuperSignal West Femto (34096)). Films were exposed in a dark room and developed following standard procedures.

Antibodies used were at 1:1000 dilution in Blocking buffer; Cleaved Caspase-3 (CST 9664), PLK1 (CST 4513), phospho-Histone H3 (S10) (Epitomics 1173-1), phospho-NDRG1 (T346) (CST5482), phospho-C-Jun (S73) (CST 3270), and 1:5000 dilution of Vinculin (CST 13901). Secondary Antibodies were used at 1:5000 dilution, Anti-rabbit IgG HRP-linked Antibody (CST 7074)

#### Retro-orbital bleeding and hematocrit determination

Mice were anesthetized using isoflurane. Blood was collected using heparinized microhematocrit capillary tubes (Fisherbrand #22-362574) through retro-orbital and was sealed with hemato-seal capillary tube sealant (Fisherbrand #02-678). First, slight pressure was applied on the eyeball to stop the bleeding. Then, Proparcanie was applied on the eye to decrease the pain after the procedure. The capillary tubes were spun at 1,500xg for 10 min. The hematocrit was calculated by dividing the measured height of red blood cell layer by the total height of the blood.

#### Intracranial orthotopic tumor cell implantation

Intracranial glioma cell injections were performed by the M.D. Anderson Intracranial Injection Core at M.D. Anderson (Dr. Fred Lang, Director^51^). Intracranial tumors were established by injection of 200,000 cells into the brains of immunocompromised female nude *Foxn1*^nu/nu^ mice ages 4-6 months, which were bred at the Experimental Radiation Oncology Breeding Core in M.D. Anderson. The animals were first bolted (a bolt is a plastic screw with a hole in the middle which is driven into the skull^51^) and allowed to recover for 2 weeks. Then, glioma cells were injected through the bolt with a Hamilton syringe. Bolting and cell injections were performed by the M.D. Anderson Intracranial Injection Fee-for-Service Core ^51^. The animals were euthanized when neurological symptoms became apparent. All animal procedures were approved by the M.D. Anderson Animal Care and Use Committee.

#### Small animal MRI and tumor volume calculation

MRI was performed in a 7T Biospec USR707/30 instrument (Bruker Biospin MRI, Billerica, MA) at the M.D. Anderson Small Animal Imaging Facility (SAIF), which is located in the same building as our mouse colony. Animals can thus be imaged serially with minimal difficulty. First, the animals are briefly maintained under deep anesthesia using Isoflurane. Body temperature was maintained with a heating blanket. Anesthetized mice were restrained using a stereotactic holder to hold their heads. Breathing was monitored and synchronized with the instrument. Routine tumor detection was done by T2-weighed imaging. First, a low resolution axial scan is taken to properly center the field. Then, a series of high resolution axial and coronal scans are recorded. The RadiAnt DICOM Viewer software was used for the image analysis.

For tumor volume calculations, we use the “ellipse” command from the tool bar to circle the tumor area in each tumor slides. The software converts the selected area from pixels to cm^2^. The axial or coronal tumor volume was calculated as both the sum of tumor volumes in each slide and the estimated tumor volume of the space in-between slides. Tumor volume for slides were calculated as the area from the RadiAnt. For coronal slicing, a thickness of 0.75 mm was used. For axial slicing, a thickness of 0.5 mm was used. For tumor volume estimations of the in-between slides, the average tumor area between the two slices was multiplied by the distance between the two slices (0.25mm for coronal slicing and 0.5mm for axial slicing). Tumor volume was then calculated as the average of coronal and axial tumor volumes.

#### Polar metabolite profiling

Culture media was removed from the plate and saved for lactate secretion determination. Adherent cells were washed once with cold PBS and aspirated completely. Four µL of ice cold 80% methanol were added to each 10 cm round plate. Plates were stored at –80 °C for 20 min. Cells were scraped from the plates on dry ice and transferred with the methanol to a pre-cooled 15mL conical tube. Next, the tube was vortexed for 5 sec and spun down at 14,000xg for 5 min at 4–8 °C to pellet the cell debris. The metabolite-containing supernatant was transferred to a clean pre-cooled 15-mL conical tube. This was then aliquoted in 1 mL fractions in 1.5 mL Eppendorf tubes and dried in the SpeedVac. The dried metabolite samples were run by tandem mass spectrometry (LC-MS/MS) via selected reaction monitoring (SRM) with polarity switching for 300 total polar metabolite targets using a 5500 QTRAP hybrid triple quadrupole mass spectrometer (SCIEX). MS was coupled to a HPLC (Shimadzu) with an amide HILIC column (Waters) run at pH=9.0 at 400 µL/min. Q3 peak areas were integrated using MultiQuant 2.1 software^52^.

#### Xenografted mouse drug treatment with POMHEX and HEX

POMHEX/POMSF were dissolved in DMSO at 200 mM and stored at –80° C. Aliquots of POMHEX DMSO stock were dissolved in PBS to achieve the desired quantity (typically 10 mg/kg, if not otherwise indicated). The total volume of 200 µL PBS (<2% DMSO) was injected intravenously or intraperitoneally. HEX was dissolved in water to a concentration of 1M; the pH adjusted to 7 with NaOH. The mouse was restrained with the Tailveiner Restrainer (Braintree Scientific #TV-150 STD). For intravenous injections, the tail was soaked in warm water for 1 min to stimulate the vasodilation, A 30G insulin syringe (BD #328431) was used for injection through lateral veins.

#### Seahorse Glycolytic Flux (ECAR) and Respiratory capacity (OCAR) determinations

The glycolytic capacity of D423 and D423 ENO1 rescued cell lines exposed to POMHEX and POMSF was assessed using XF Glycolysis Stress Test Kit (Seahorse Bioscience) according to the manufacturer protocol. Briefly: 40 × 104 cells/well were plated on a 96-well Seahorse microplate 24 h before the experiment. (DMEM supplemented with 10% FBS and 1 % Pen-Strep). The medium was replaced 1 h before the experiment with Glycolysis Stress Test Assay Medium supplemented with 2 mM L-glutamine. Cells were then incubated at 37° C degree (low CO_2_) for 1h. The Seahorse cartridge was hydrated for 24 h in calibrating solution (Seahorse Bioscience) followed by loading with POMHEX or POMSF – PORT A (concentration range from 1 µM to 0.062 µM), glucose (10 mM) – PORT B and oligomycin (2.5 µM) – PORT C. The extracellular acidification rate (ECAR) was measured using the Seahorse XF analyzer (Seahorse Bioscience). POMHEX and POMSF were added for 150 min before glycolytic capacity of the cells was assessed. After measurement, the cells were fixed for 10 min in 10% NBF followed by 15 min incubation in Hoechst 33332. The fluorescence of each well was quantified using Tecan Infinite M200PRO Plate Reader.

### Pharmacokinetic quantification of POMHEX and its metabolites

POMHEX: 25 µL of plasma sample, including the analytical standards and quality controls (QCs), were mixed with 200 µL of ACN. After vortex and centrifugation, 100 µL of the supernatant was mixed with 100 µL of water for LC-MS/MS analysis. The same procedure was followed for the preparation of cell lysate samples.

HemiPOMHEX and HEX: 50 µL of plasma sample including the standards and QCs were diluted with 500 µL of 10 mM aqueous NH_4_HCO_3_ containing Bn-HEX (0.015 µM, internal standard). This was then loaded onto a Waters Oasis MAX SPE cartridge (30 mg) which was pre-conditioned with methanol (1 mL) and water (1 mL). The cartridge was washed with water (1 mL) and methanol (1 mL). Analytes were derivatized on the cartridge with 200 µL of 2.0 M solution of trimethylsilyl-diazomethane in hexane (TMS-DAM, Sigma Aldrich) for 1 h to obtain TMS-derivatized Hemi-POM-HEX and HEX. The derivatized analytes in the cartridge were eluted with methanol (1 mL), evaporated to dryness at 40 °C under a stream of nitrogen, and reconstituted with 150 µL of 0.1% FA in water/ ACN (2:1) for LC-MS/MS analysis. The same procedure was followed for the preparation of cell lysate samples. Samples were analyzed on an Agilent 1290 Infinity Binary LC/HTC Injector coupled to a Sciex 5500 Triple Quadrupole Mass Spectrometer operated in positive mode. The mass spec source conditions were set for all the analytes as the following: ion spray voltage (5500 volts), curtain gas (35), collision gas (8), gas temperature (500° C), ion source gas 1 (40), ion source gas 2 (60), DP (156), EP (10), CE (27), CXP (10). The LC mobile phase A was 0.1% acetic acid-water and B was 0.1% acetic acid-acetonitrile. The LC flow rate was 0.5 mL/min. The injection volume was 2 µL. The column temperature was set to 40° C. POMHEX was separated using a Supelco Ascentis fused-core C18 (2.7 µm, 2.1 × 20 mm) column (Sigma-Aldrich, St. Louis, MO) and detected by a multiple reaction monitoring transition (m/z 424.1>280.1). The LC gradient was 25% B (0-0.3 min), 25-95% B (0.3-1.3 min), 95% B (1.3–1.7 min), 95-22% (1.7–1.71 min), 22% B (1.71–2.0 min). Under these conditions, the retention time of POMHEX was 1.08 min. The method was validated with an analytical range of 5 – 1000 ng/ml for POMHEX in untreated CD-1 mouse plasma. The acquired data was analyzed using Sciex MutiQuan 3.0.2 software. The TMS-derivatized HemiPOMHEX and HEX were separated using a Phenomenex Luna C8 (3 µm, 2.0 × 50 mm) column (Torrance, CA). The multiple reaction monitoring transition was 360.4>330.1 (TMS-HemiPOMHEX), 238.1>207.0 (TMS-HEX), and 313.9>91.1 (TMS-Bn-HEX, IS). The LC gradient was 5% B (0-0.3 min), 5-95% B (0.3-1.5 min), 95% B (1.5-2 min), 95-22% (2-2.1 min), 22% B (2.1-2.3 min). Under these conditions, the retention times were 0.97 min (TMS-HEX), 1.30 min (TMS-Hemi-POM-HEX), and 1.26 min (TMS-Bn-HEX, IS). The method was validated with an analytical range of 0.1 – 5 µM for both Hemi-POMHEX and HEX in untreated CD-1 mouse plasma.

#### Lactate secretion determinations by NMR

Cell culture media was spun down at 14,000 g for 10 min at 4°C. The supernatant was transferred to a clean, pre-cooled 50 mL conical tube. Then, 40 mL of pre-cooled methanol was added to the 10 mL of supernatant to make a final 80% (vol/vol) methanol solution. The mix was gently shaken and incubated overnight at –80°C. Next the mixture was spun down at 14,000 g for 10 min (4–8 °C). The supernatant was transferred to a new 50 mL conical tube and dried in the SpeedVac. The dried metabolites were re-suspended with 400uL of D2O and 50uL of D2O with TMS (ALDRICH #450510) for NMR analysis. Nuclear Magnetic Resonance (NMR) analysis was performed at the MD Anderson Core Facility, supported in part by the Cancer Center Support Grant from the National Cancer Institute (CA16672). A 500 MHz Bruker Avance III HD NMR equipped with a Prodigy BBO cryoprobe instrument was used. Conditioned media was supplemented with 10% D2O for signal lock and 3-(trimethylsilyl)propionic-2,2,3,3-d4 acid standard. Lactate exhibits a highly characteristic, strong doublet peak at 1.4 ppm in the 1H spectrum, which was quantified by integration relative to the 3-(trimethylsilyl)propionic-2,2,3,3-d4 acid standard at 0 ppm.

#### ENO2:HEX crystal structure

Apo crystals of Human Enolase 2 were prepared using the hanging drop vapor diffusion method at 20 °C, suspending a drop containing 0.5 μL 9.1 mg mL-1 Enolase 2 and 0.5 μL reservoir solution above a 500 μL reservoir containing 200 mM ammonium acetate, 100 mM Bis-Tris and 18-22% (w/v) PEG 3350. Streak seeding in the same solution conditions reproducibly produced larger crystals. The crystals were soaked overnight in 1 μL drops containing 100 mM Bis-Tis, 200 mM ammonium acetate, 32% (w/v) PEG 3350 at pH 65, supplemented with 2 mM HEX compound, prior to flash freezing in liquid nitrogen. A X-ray diffraction dataset was collected at 100 K using the Advanced Light Source Beamline 8.3.1 equipped with ADSC Q315r detector. 360 images with 1° oscillation were collected, using 1.116 Å wavelength. The diffraction images were indexed and integrated using iMOSFLM ^53^ and scaled using AIMLESS^54^. The X-ray structure was solved by molecular replacement with a human Enolase 2 homodimer (PDB code 4ZCW) as the search model. The structure was iteratively refined using Coot^55^ and phenix.refine^56,57^.

### Statistical Analysis

Statistical significance was analyzed using the unpaired, two-tailed, t-test function of Microsoft Excel or Prism GraphPad. The Welsh correction was used for unequal variance and the Bonferroni correction was used for multiple hypothesis testing.

## Acknowledgments

We thank James Holton and George Meigs for their assistance with X-ray diffraction data collection. We thank Kumar Kalurachchi and John McMurray for assistance with NMR. We thank Steven Millward, Seth Gammon for critical comments and suggestions. We thank Min Yuan and Susanne Breitkopf for help with the mass spectrometry experiments. We thank Hannah Butterfield, Lexiah Jacobs, and Pornpa Suriyamongkol for technical assistance. We thank Christopher Halbrook and Costas Lyssiotis for helpful discussions on metabolomics. We thank Keith Michel, Charles Kingsley, Vivien Tran and the rest of the SAIF team for expert assistance with MRI imaging; Verlene Henry, Joy Gumin and Fred Lang for GSC’s and intracranial cell implantations. Pharmaceutical Chemistry Facility at MD Anderson supported by NIH/NCI under award number P30CA016672 (R.A.D.). Financial support was provided by CPRIT RP140612 (R.A.D.) and NIH CDP SPORE P50CA127001-07 (F.L.M.). ACS RSG-15-145-01-CDD (F.L.M),NCCN YIA170032 (F.L.M. Young Investigator Award). This work was also supported by The University of Texas MD Anderson Cancer Center Institutional Research Grant (IRG) Program. This work is dedicated to the memory of Dr. John Stuart McMurray, an expert in phosphonate pro-drug synthesis, without whom this would not have been possible.

## ACCESSION CODES

X-ray structures of ENO2 bound with HEX (5IDZ) were deposited in PDB.

